# Large-scale rewiring in a yeast hybrid

**DOI:** 10.1101/100263

**Authors:** Rebecca H. Herbst, Dana Bar-Zvi, Sharon Reikhav, Ilya Soifer, Michal Breker, Ghil Jona, Eyal Shimoni, Maya Schuldiner, Avraham Levy, Naama Barkai

## Abstract

The merging of genomes in inter-specific hybrids can result in novel phenotypes, including increased growth rate and biomass yield, a phenomenon known as heterosis. We describe a budding yeast hybrid that grows faster than its parents under different environments. Phenotypically, the hybrid progresses more rapidly through cell cycle checkpoints, relieves the repression of respiration in fast growing conditions, does not slow down its growth when presented with ethanol stress, and shows increasing signs of DNA damage. A systematic genetic screen identified hundreds of alleles affecting hybrid growth whose identity vastly differed between the hybrid and its parent and between growth conditions. This large-scale rewiring of allele effects suggests that despite showing clear heterosis, the hybrid is perturbed in multiple regulatory processes. We discuss the possibility that incompatibilities contribute to hybrid vigor by perturbing safeguard mechanisms that limit growth in the parental background.

## Introduction

Hybrids between related species or strains often display traits that are superior to their parents, in particular in relation to growth vigor. This phenomenon, known as heterosis, has been observed in all eukaryotic kingdoms: plants, animals and fungi. Hybrids and heterosis has fascinated evolutionary biologists since Darwin^1,2^ and continue to fascinate modern day geneticists. Heterosis is often viewed as the opposite of hybrid incompatibility, the more expected clash between genomes^3^.

Hybridization plays an important role in the emergence of new species. Hybrid vigor can give an advantage to hybrids in certain niches, and hybrid incompatibilities can secure reproductive isolation^4–8^. In addition to its evolutionary importance, hybrid vigor has been extensively exploited for increasing productivity in agriculture^9,10^. Yet, despite extensive research, the underlying mechanism of hybrid vigor remains elusive^3^. The common view, “the dominance model” describes heterosis as the opposite of inbreeding depression, i.e. in the hybrid state, deleterious mutations specific to one parent, are complemented by the wildtype dominant allele of the other parent^11,12^. A second class of models^3^ attributes heterosis to overdominance or epistasis effects, in which interactions between alleles of the two parents, either coding for the same gene or for different genes, emerge in the hybrid and lead to its superior performance. Studies of heterosis in different plant species provided support for both the dominance and overdominance model by defining specific alleles that contribute to heterosis. The number of alleles with a known mechanism remains limited, and the extent to which heterotic effects are distributed across the genome is not known.

To gain a broader view on the organization of the hybrid’s regulatory processes, genome-wide profiling approaches were applied. These studies revealed large-scale differences between hybrids and their parents in gene expression^13,14^, nucleosome positioning^15^, and other genomic features^16,17^. The unique expression pattern in the hybrid was attributed mainly to differences between the parents: most genes were expressed at a level that was intermediate between the two parents. An additional fraction of genes showed a distinct expression, being expressed more, or less than both parents, likely resulting from novel cis-trans interactions that emerged in the hybrid due to the mixing of the two parental genomes. Gene expression rewiring could affect regulatory mechanisms and thereby contribute to the emergence of novel phenotypes, including heterosis. However, whether and how changes in gene expression translate into growth effects remains unclear.

The widespread differences in gene expression between the hybrid and its parents raised the question of how heterotic loci are distributed across the genome. Can we attribute heterosis to a small number of genes, or is it the results of multiple effects distributed across many alleles? Are heterotic effects confined to specific functional groups, and if so, which processes are most prone to such effects? To examine this, we turned to budding yeast, where systematic genetic screens are more easily performed. A recent study reported that heterosis is relatively common in budding yeast, being observed in 35% out of the 120 intra-specific crosses between different strains analyzed^18^. Inter-specific hybrids are also widespread in domesticated strains used for the making of alcoholic beverages, bread and biofuel^19^, but they have received less attention in heterosis research.

Budding yeast hybrids are easily generated in the laboratory, and in our initial studies we observed a particular hybrid between *Saccharomyces cerevisiae* and *Saccharomyces paradoxus* that grew faster than both its (fast growing) diploid parents under multiple conditions. Using this model, we examine systematically the contribution of all viable *S. cerevisiae* alleles to hybrid growth. We identified hundreds of alleles that contribute to hybrid growth, but show no effect in the parental background. Conversely, a large number of alleles showed a dosage effect in the parental background, but did not contribute to hybrid growth. The multiplicity of allele effects that were specific to the hybrid, together with their functional associations, suggest that the hybrid experiences not only growth heterosis but also incompatibilities that dysregulate evolved regulatory mechanisms, a notion that is supported by our phenotypic analysis. If these incompatibilities target safeguard processes that limit growth in the parental background, then they could directly contribute to growth heterosis, as we discuss.

## Results

### A budding yeast hybrid showing growth heterosis

We generated hybrids by mating haploids of *S. cerevisiae* and *S. paradoxus*, two closely-related *sensu-stricto* species that express largely the same set of genes at preserved ontology, with 90% and 80% sequence identity in coding or inter-genic regions, respectively^20^. The hybrids are limited in meiosis, but can propagate vegetatively without signs of genetic stability or aneuploidy. We previously used this hybrid for studying regulatory divergence^13^. Here, we describe its phenotypic properties and use it to systematically evaluate how heterotic alleles, defined here as alleles that contribute to the growth of the hybrid but do not have an effect in the parental background, are distributed across the genome.

When provided rich media (SD), both diploid parents grow with a division time of ~90 minutes, typical of rapidly growing strains. Still, the hybrid grows ~20% faster than both diploid parents (Fig. 1A). Growth heterosis was observed also in other conditions, including high temperature, presence of high ethanol and low-Pi (Fig. 1A). We used live-cell microscopy to quantify the duration of the different cell cycle phases (Fig. 1B, Movies S1-3). The two parents regulate their cell cycle differently: as described before, *S. cerevisiae* cells have short G2 phase, thereby generating small daughter cells that extend their G1 phase to retrieve their mother’s size. *S. paradoxus*, on the other hand, regulate its size by extending the G2 phase, followed by a short G1^21^. We find that the hybrid G2 phase was as short as in *S. cerevisiae*, yet it did not extend its daughter’s G1, which was as short as in *S. paradoxus*, (Figure 1C, S1).

**Fig. 1.**
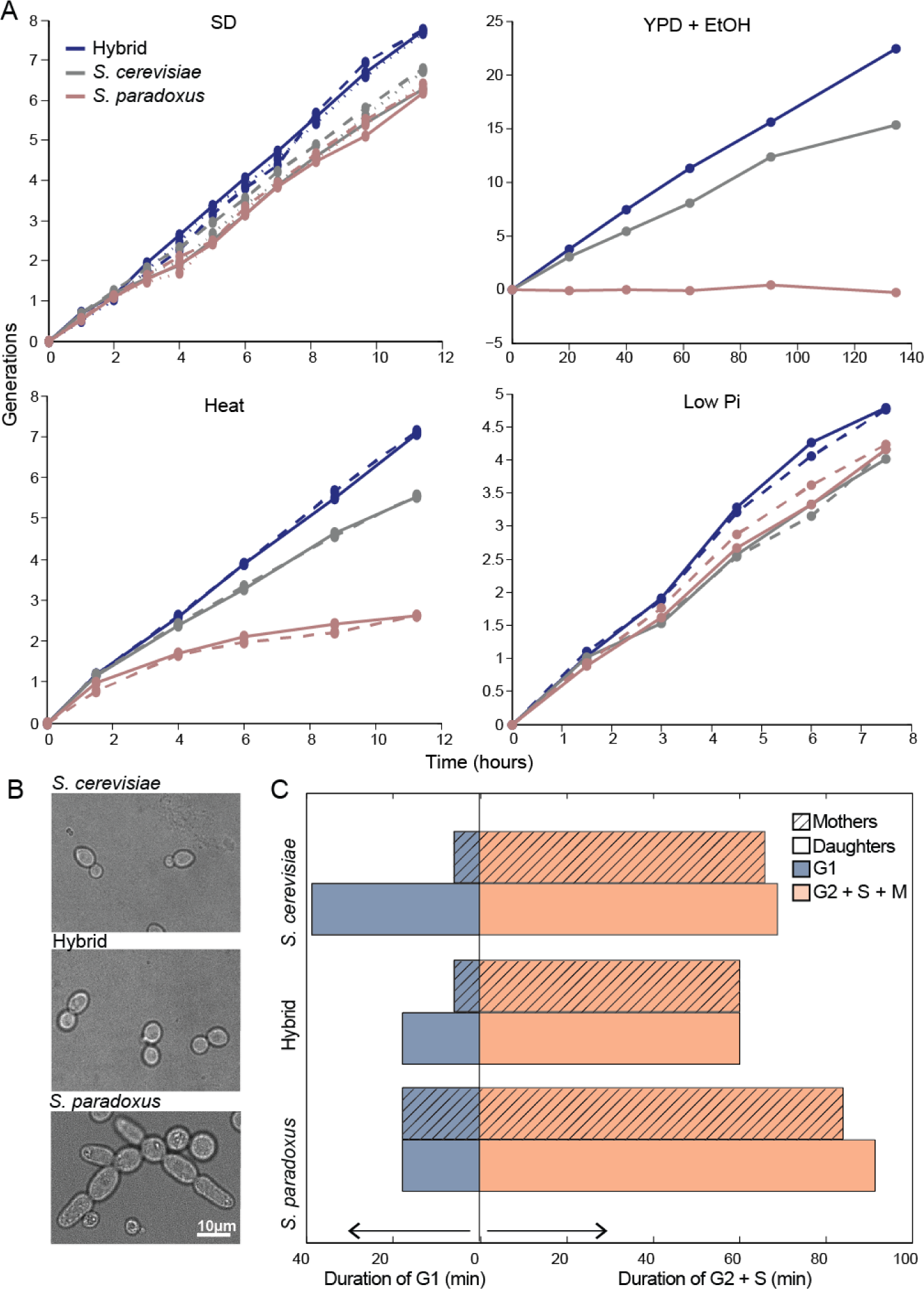
Growth heterosis in a yeast hybrid. **(A)** *Growth heterosis in the yeast hybrid in rich and stress media*: cells were diluted into rich (SD, YPD) or stress media (YPD+7.5% ethanol, YPD in 37° C, 0.2mM Pi), and doubling time was measured by following their optical density (OD). Cultures were diluted periodically to maintain the cells in logarithmic phase for the duration of the experiment. Different repeats indicate biological replicas. **(B)** *Growth pattern of the hybrid and its diploid parents presented with rich (SD) media* **(C)** *Perturbed cell-cycle delays in hybrid*: Shown are the median cell-cycle times of the unbudded (G1) and budded (S+G2+M) hybrid and its diploid parents. Mother and daughters are shown separately. Data extracted from live-cell microscopy (see Movies S1-3, and time distributions in fig. S1).

Sustained rapid growth entails a more efficient production of biomass. Biomass and energy production are regulated by the routing of carbon through central carbon metabolism. We previously noted that respiration gene expression is higher in the hybrid relative to its parents, even when grown in glucose (Fig. S13 in ^13^, Fig. 2A), which was surprising, as budding yeast represses respiration genes and does not respire in the presence of glucose. We therefore asked whether glucose repression is reduced in the hybrid, enabling a more efficient energy generation through respiration. Measuring oxygen usage along the growth curve confirmed that hybrids consumed oxygen at a high rate throughout the growth curve, even when glucose was abundant, in contrast to *S. cerevisiae* and *S. paradoxus* where oxygen consumption was significantly lower when glucose was present (Fig. 2B, S2A-B). Consistently, hybrid mitochondria were significantly larger and contained more cristae compared to *S. cerevisiae* and *S. paradoxus*, as visualized by electron microscopy (EM) (Fig. 2C-D, S2C). Finally, heterosis was reduced upon the addition of the respiration blocker Antimycin A to the level of the best parent (Fig. 2E, S2D). Together, our results suggest that reduced glucose repression in the hybrid enables it to respire even in the presence of glucose.

**Fig. 2.**
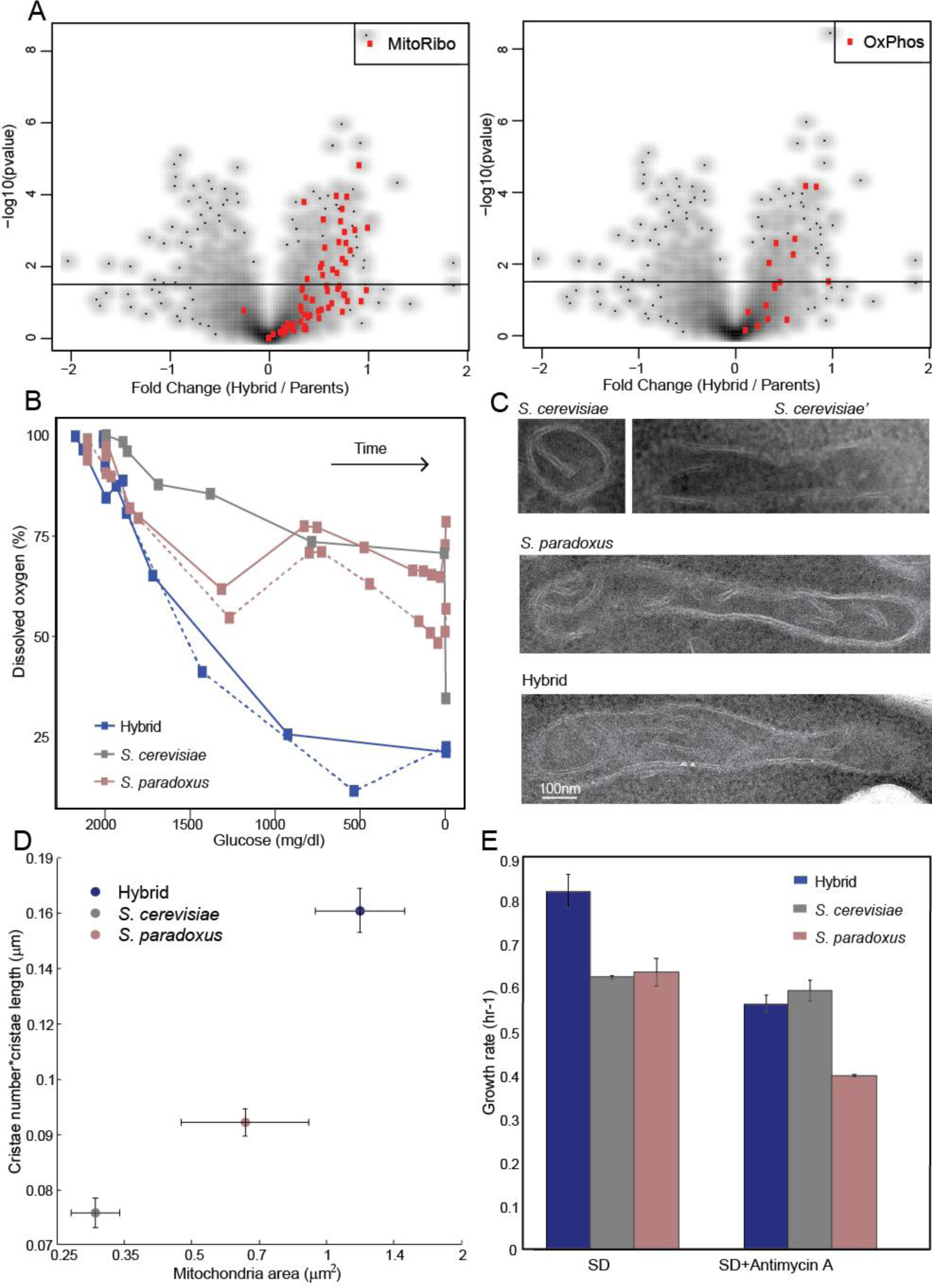
Perturbed glucose-dependent respiration repression in the hybrid. **(A)** *Reduced glucose-repression through upregulation of respiration gene expression in the hybrid*: each point represents a gene from the indicated group. Fold expression changes were calculated between the hybrid and the mean expression of *S. cerevisiae* and *S. paradoxus*. See Table S1 for gene names and expression values. Data from^13^. **(B)** *The hybrid consumes oxygen in the presence of glucose, but not its diploid parents*: Shown are measurements of glucose and dissolved oxygen for cultures incubated into glucose-containing rich media (SD). Measurements were made during batch growth within a fermentor. **(C-D)** *Hybrid mitochondria are larger and contain more cristae*: (C) Representative EM images of mitochondria in the hybrid and its diploid parents grown in SD. (D) Quantification of mitochondria area and cristae of the hybrid and the diploid parents, from EM images. **(E)** *Heterosis is lost when respiration is inhibited*: Growth rates are shown for cells growing in SD in the presence or absence of the respiration inhibitor Antimycin A.

As an additional test, we examined the instances of DNA damage in the hybrid, by following Rad52-GFP, a protein that localizes to foci of DNA double-strand breaks^22^. Notably, the frequency of cells with Rad52-GFP foci increased ~two folds in the hybrid compared to either parent (Fig. 3A). Further, we noted that in the hybrid, a group of cytosolic chaperones localized into punctate structures, a known marker for DNA damage stress (Fig. 3B, table S2).

**Fig. 3.**
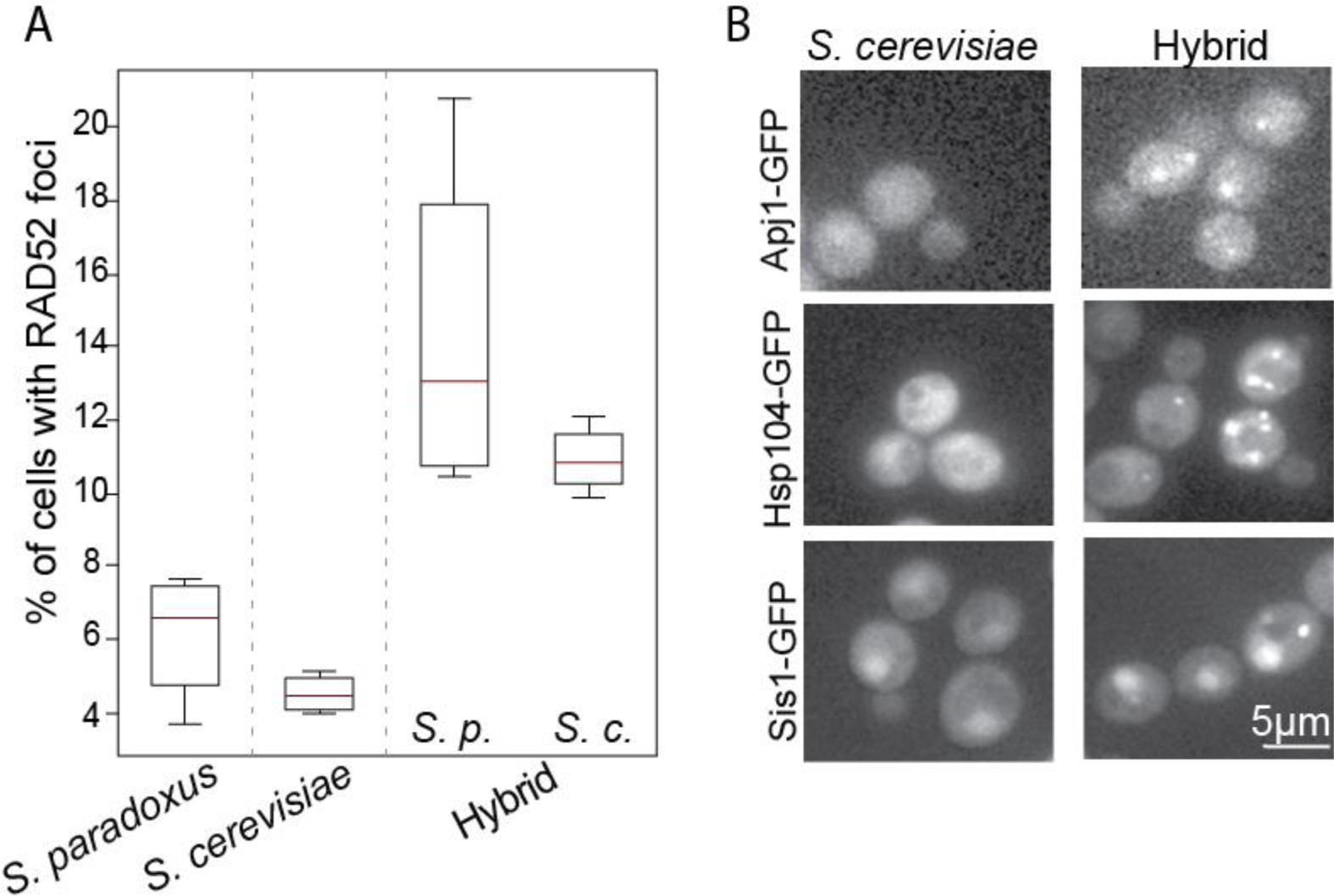
Increased DNA damage in the hybrid. **(A)** *Increased formation of foci indicative of DNA damage in hybrid*: fraction of cells showing *RAD52-GFP* foci in the hybrids and its diploid parents grown in SD. For the hybrid, the results are shown when tagging either the *S. cerevisiae* or *S. paradoxus* allele. **(B)** *The hybrid shows signs of DNA damage stress through protein localization*: chaperones that are known to localize into punctates during DNA damage are observed in the hybrid (Table S2).

### A genome-wide genetic screen detects hundreds of alleles contributing specifically to hybrid growth in an environment-dependent manner

Classical models of heterosis, including dominance, overdominance or dosage models, attribute the hybrid’s superior performance to the action of specific alleles. We reasoned that, working with budding yeast, we can systematically screen for heterotic alleles, defined here as alleles which contribute to the hybrid’s growth but do not show a dosage effect in the parental background. This general definition includes alleles that function through dominance, dosage, overdominance, epistasis or more complicated (e.g., cis and trans) effects.

Our screen is based on the availability of a deletion library, corresponding to all non-essential *S. cerevisiae* genes. Starting from this library, we generated a library of hemizygote hybrids that lack a specific *S. cerevisiae* allele but contain the corresponding *S. paradoxus* allele, and a control library containing hemizygote *S. cerevisiae* diploids (Fig. 4A). Two independent libraries were generated for each genetic background.

**Fig. 4.**
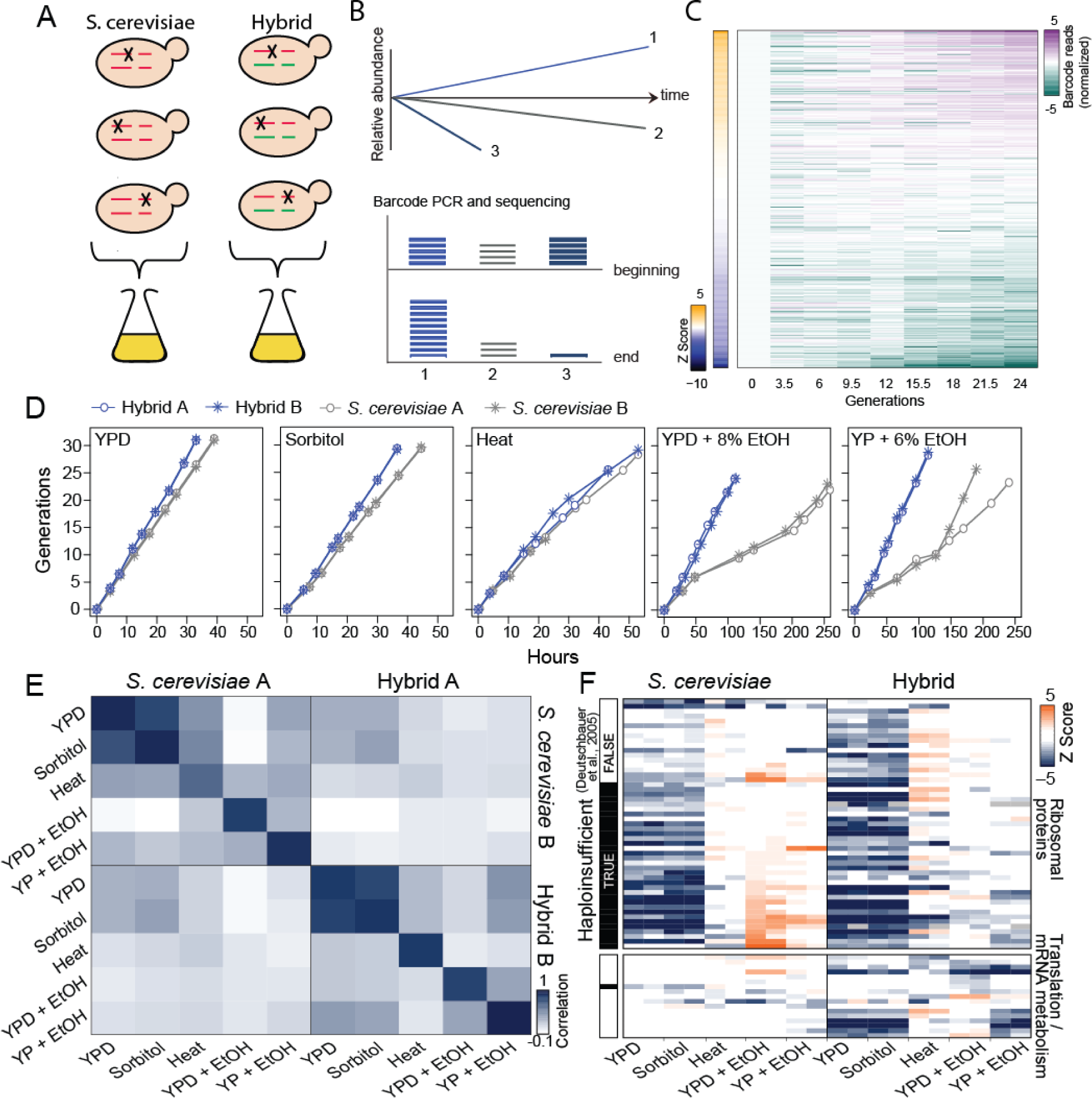
Genome-wide screen for alleles contributing to hybrid growth. **(A-B)** *Screen for S. cerevisiae alleles contributing to hybrid growth*: A hemizygote hybrid library was constructed by mating *S. paradoxus* with a library of *S. cerevisiae* strains, each of which was deleted of one *S. cerevisiae* gene. Each strain was labeled by a specific barcode, enabling the measurements of relative strain abundances within a co-growing pool using sequencing. **(C)** *Change in abundance of individual hemizygote strains*: a pool of hybrid hemizygotes was grown in YPD+8% ethanol. Shown are the (log) normalized barcode reads of 100 strains showing the most pronounced increase or decrease in frequency, together with 200 control strains that maintained their abundance throughout the experiment. The Z-score of the calculated growth rate for each strain is shown in the sidebar. **(D)** *Growth curves of hemizygote hybrid and S. cerevisiae pools:* The two hemizygote pools were kept for ~30 generations in logarithmic phase through subsequent dilution, and sampled every three generations. Two independent biological replicates are shown. **(E)** *Correlation between growth rates of hemizygote strains*: Note the reproducibility between the two independent repeats, contrasting the low-correlation when comparing different conditions or backgrounds. **(F)** *Hybrid-specific dosage sensitivity*: shown are z-scores of growth rates of strains hemizygote for genes coding for ribosomal proteins and mRNA metabolism, as indicated, that show reproducible effect in at least one condition (Z Score < -1.5 ; see Table S5 for gene list). Missing values are shown in black. Sidebar marks previously defined haploinsufficient genes^24^.

As described previously, the deletion library was specifically designed to enable sensitive measurement of the growth rate of individual strains, while growing all strains in one pool. This approach has the advantage that all strains are exposed to the same environment, making a comparison between the individual strains more controlled. Specifically, each strain in the library is marked by a specific sequence barcode that is flanked by a common sequence, so that high-throughput sequencing can be used to quantify the relative abundance of each strain within the growing pool (Fig. 4B-C, see Methods, Ref^23^.). Temporal changes in a strain’s abundance during pool growth then report on its growth rate relative to the pool’s average: slow growing strains will be gradually outcompeted, while the fast growing ones will become increasingly more abundant. Note that the measured growth rate of the pool well approximate the wild-type growth rate, as the majority of mutants do not show a growth defect at any given condition^23^.

We considered five growth conditions (YPD, YPD + Sorbitol, YPD at 37°C, YPD + 8% ethanol, YP + 6% ethanol; Fig. 4D, Table S3-4). In each condition, the pools were maintained in log-phase throughout the experiment, to ensure constant conditions and limit possible effects of nutrient depletion or secretion. In total, 808 alleles reduced growth in at least one condition or genetic background (*Z* Score < -1.5 in both biological replicates, fig. S3A). These genes were classified into functional groups based on databases and literature (fig. S3B, Table S5). Inferred growth rates were highly correlated between two replicates corresponding to two independently pooled libraries.

In contrast to the high reproducibility of effects between independent repeats, different conditions showed little similarity (Fig. 4E). For example, stress conditions increased dosage sensitivity to genes involved in peroxisome function, cell wall formation/breakdown, or protein and lipid modifications (fig. S3C). Most notably, while previously annotated haploinsufficient genes in *S. cerevisiae^24^* such as ribosome components, were reproducibly identified as dosage sensitive in rich media, these strains showed no effect on the slower growth conditions (Fig. 4F, S3D). Hence, loss of one allele could range from being deleterious to even providing a growth advantage depending on the growth condition.

Surprisingly, perhaps, the set of genes sensitive to hemizygosity in the hybrid greatly differed from that of its *S. cerevisiae* parent, even within the same growth condition (Fig. 4E-F). In fact, the low correlation between the different growth conditions in the hybrid and *S. cerevisiae* was comparable to the low correlation between the hybrid and the parent in the same condition. Therefore, hundreds of *S. cerevisiae* alleles contribute to hybrid growth but not to *S. cerevisiae* growth. These alleles are distributed across the genome, being involved in multiple processes and pathways (Fig. 4F, S3B-C).

### Heterotic alleles consistent with hybrid’s phenotypes

We observed a correspondence between allele-specific sensitivity and the hybrid phenotypes noted above. First, the hybrid shows increased sensitivity to alleles involved in the G1/S transition (Fig. 5A) and to alleles whose deletion increases cell size, irrespectively of their functional association (hypergeometric p-value: Hybrid < 10^−10^, *S. cerevisiae*: 0.65, Fig. 5B). This increased sensitivity may be explained by the shorter duration of this cell cycle phase in the hybrid compared to *S. cerevisiae*: as G1/S transition is a major checkpoint where nutrition status and cell-size are monitored, its shorter duration may limit the checkpoint capacity to correct for size perturbations.

**Fig. 5.**
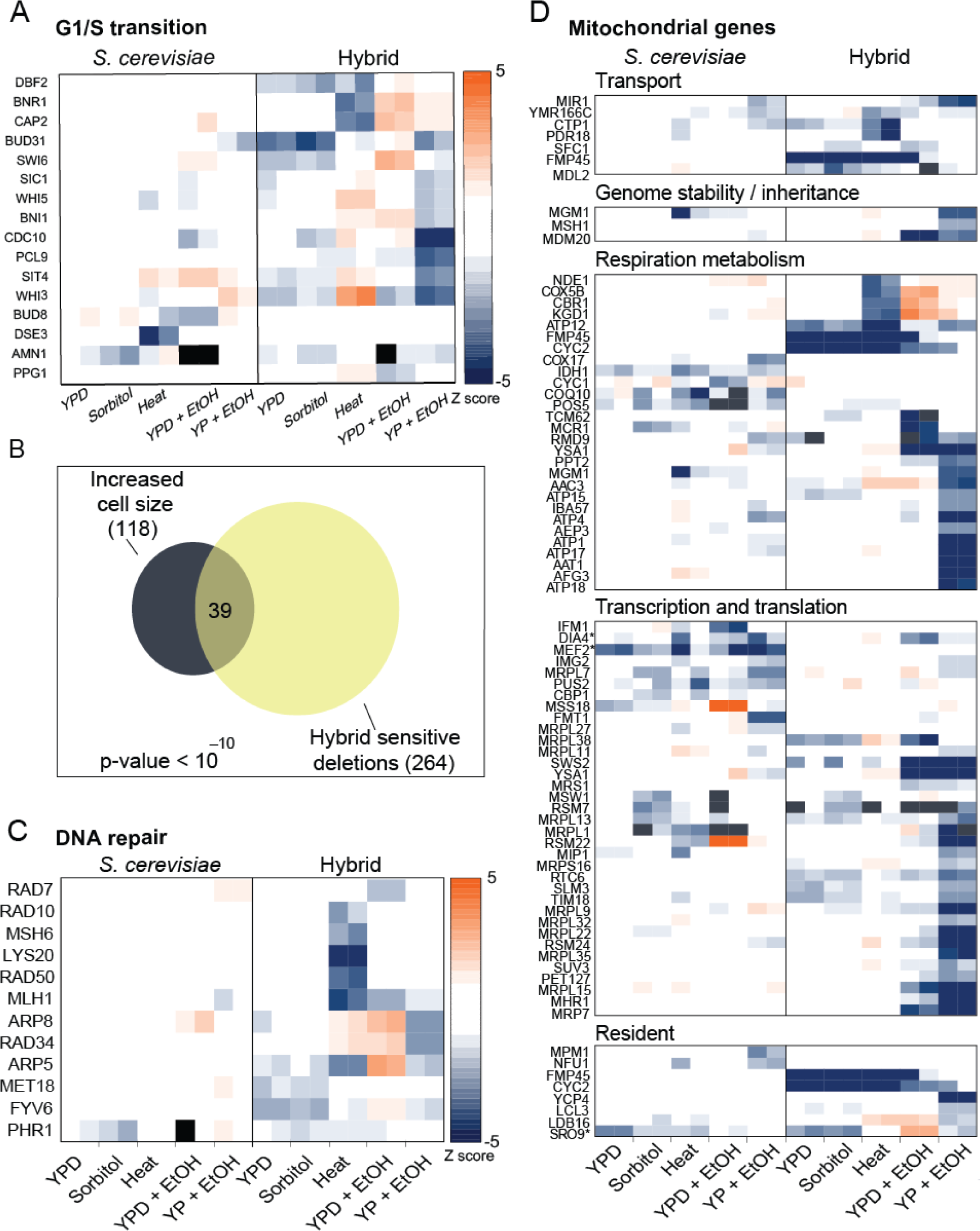
Gene sensitivity results indicate perturbation in several pathways. **(A)** The *hybrid shows increased sensitivity to cell-cycle genes*: shown are z-scores of growth rates of strains hemizygote for genes coding for G1/S transition that show reproducible effect in at least one condition. Missing values are shown in black. **(B)** The *hybrid shows increased sensitivity to genes increasing cell size*: Genes reported to increase cell size^39^ are highly enriched within the set of genes showing hybrid-specific dosage sensitivity in at least one of the conditions (hypergeometric test). This increased sensitivity is independent of the functional association of these genes. **(C)** The *hybrid shows high sensitivity to DNA damage genes*: Same as Fig. 5A for the indicated strains. **(D)** *Hybrid hemizygote strains show high sensitivity to mitochondrial genes*: Same as Fig. 5A for the indicated genes, which code for proteins that localize to the mitochondria and are either nuclearly or mitochondrial encoded.

Also connected to cell cycle progression, the hybrid showed an increased sensitivity to DNA repair genes (Fig. 5C). This, together with the phenotypic results of increased presence of DNA damage markers in the hybrid (Fig. 3A-B), may suggest suboptimal performance of mechanisms maintaining genome integrity.

Finally, consistent with the reduced glucose repression seen in the hybrid, the hybrid showed increase sensitivity to mitochondrial genes compared to the *S. cerevisiae* parent (numerical p-value < 10^−2^). This was particularly pronounced during growth on ethanol, but observed also on glucose (Fig. 5D).

### The hybrid escapes a programmed cell cycle slow-down under severe ethanol stress

The largest difference in the pattern of allelic sensitivity between the hybrid and *S. cerevisiae* was observed under conditions of high ethanol stress. Under this condition, a reproducible minority of hemizygote strains overtook the population in the two *S. cerevisiae* replicates (Fig. 6A-B, S4A-B, Table S6). In contrast, the hybrid showed the typical pattern of allele sensitivity as seen in other conditions. This differential pattern of effects was also reflected in the growth of the pools (cf. Fig. 1A, 4D): while the hemizygote hybrids maintained steady growth throughout the experiment, similarly to all other conditions tested, growth of the hemizygote *S. cerevisiae* pool was initially rapid, then slowed down, and became rapid again after ~10 generations. The rapid hybrid growth in high ethanol is especially striking, considering that the *S. paradoxus* parent fails to grow in this condition (Fig. 1A).

**Fig. 6.**
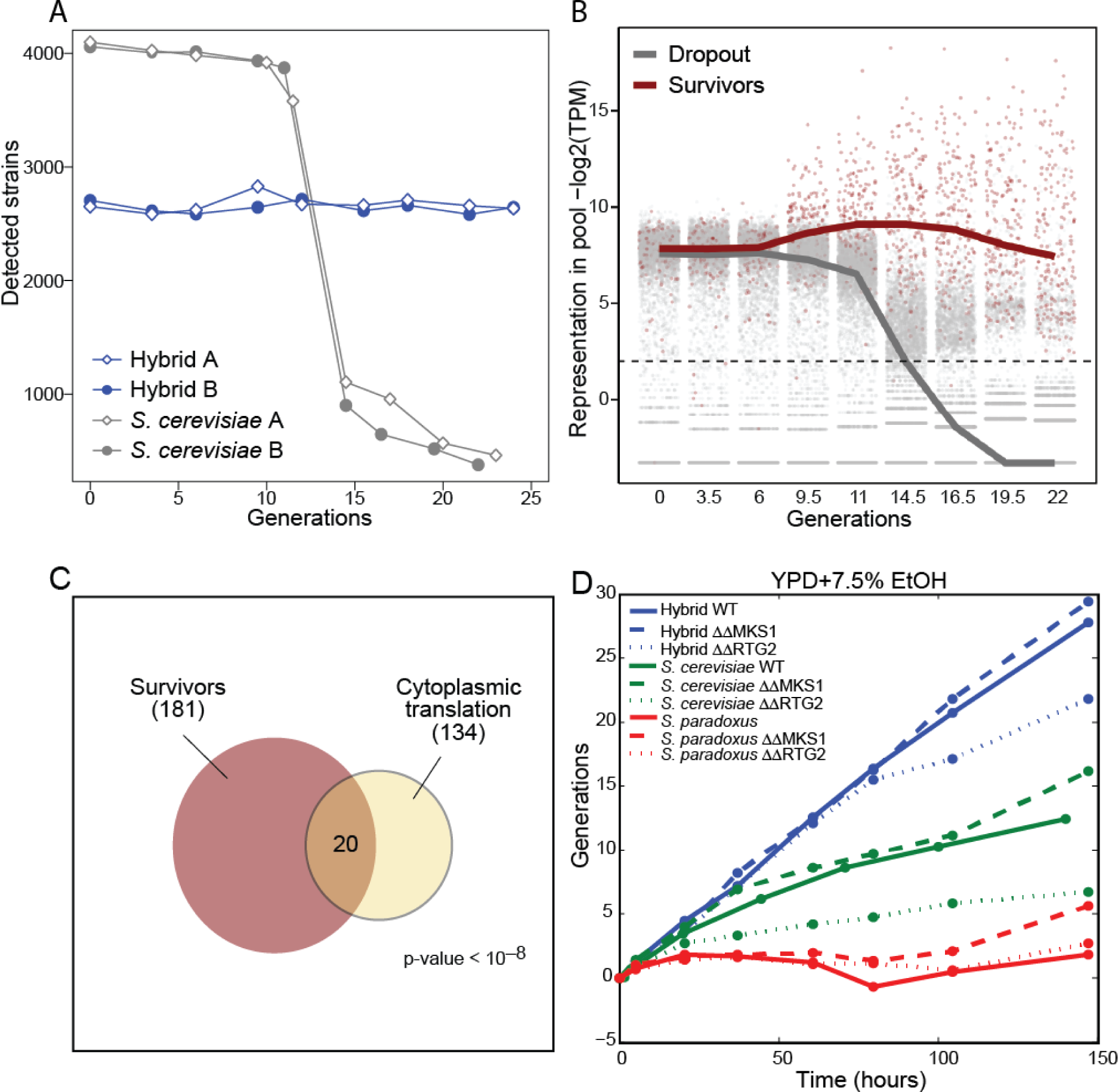
Hybrid does not slow-down its growth during ethanol stress. **(A-B)** *A minority of hemizygote strains overtakes the S. cerevisiae pool when subject to ethanol stress*: the number of strains that were reliably identified (>30 reads) when sequencing the hemizygote hybrid or *S. cerevisiae* pools at different time points in YPD + 8% ethanol medium is shown (A) and the relative representation of each strain within the *S. cerevisiae* pool is shown in (B). Red points represent the strains that were abundant (log2(TPM+1) > 2) at the last two time points in both *S. cerevisiae* experiments. **(C)** *Strains deleted for genes coding for ribosomal proteins are enriched in the survivor pool*: Shown is the hypergeometric p-value for enrichment of ribosomal genes. **(D)** *Increasing retrograde signaling improves hybrid growth under ethanol stress*: Cultures were diluted periodically to maintain the cells in logarithmic growth. MKS1 is a negative regulator, whereas RTG2 is an activator of retrograde pathway. ΔΔ refers to strains deleted of both alleles of the gene.

The hybrid therefore maintains a stable growth in high ethanol, while the *S. cerevisiae* diploids slowdown their growth following some period. Notably, this slowdown can be overcome by decreased expression of individual genes. This unique dosage response suggests that the growth slow-down is an active and adapted strategy and not a passive reaction to unavoidable toxicity. In support of that, strains that are maladapted in rich conditions were enriched amongst the surviving hemizygote diploids (Fig. 4F, 6C).

To try and identify the basis of this increased ethanol resistance in the hybrid, we examined the pattern of allelic sensitivity. The hybrid showed an increased dependency on retrograde signaling (fig. S4C). This response is triggered by damaged mitochondria to induce nuclear-encoded protecting mechanisms^25^. Induction of this pathway within the hybrid could render the hybrid more stress resistant. In support of that, over-activating the retrograde pathway increased growth under ethanol stress for both the hybrid and its *S. cerevisiae* diploid parent, although accounting for only a fraction of hybrid growth vigor (Fig. 6D, S4D).

## Discussion

### The genetic control of heterosis

We describe a systematic analysis of hybrid growth in the absence of a specific allele from the *S. cerevisiae* parent. The advantage of this screen is that it enables testing the effect of a genetic locus in the relevant hybrid background rather than in segregating populations or after introgression in a non-hybrid background (e.g. recombinant inbred or introgression lines). Although testing only one set of parental allele, our analysis provided us with a comprehensive characterization of alleles that contribute to hybrid growth.

We initially expected that dosage sensitivity would be largely similar between the parental and hybrid backgrounds, allowing us to identify a limited number of alleles that contribute to heterosis through dominance or partial dominance effects. In striking contrast to these expectations, hundreds of loci identified to be sensitive to hemizygosity in the hybrid greatly differed from those loci in its *S. cerevisiae* parent, even within the same growth condition (Fig. 4E-F). Furthermore, the set of hemizygote-sensitive genes also reproducibly varied between conditions. While this could indicate that dominance effects are abundant in this hybrid, we find this to be unlikely for several reasons. First, the number of alleles that had an effect in the hybrid but not in the parent was equivalent to the number of alleles that affected the parent but not the hybrid. Second, the differences in allele effects between the hybrid and its parent were similar to the differences in allele effects between *S. cerevisiae* cells growing in different conditions. Finally, both parents grow at a rate that is characteristics of fast growing strains, suggesting that they do not contain a large number of deleterious alleles. We therefore favor the alternative possibility that most effects represent differential dosage sensitivity between the hybrid and its parent. Notably, studies in rice^26^ showed that most loci involved in heterosis were not dominant, but rather partial dominant, which is consistent with dosage sensitivity. Recent studies on the *SINGLE FLOWER TRUSS* locus in tomato also support the importance of dosage optimality in heterosis^27,28^.

### The Dysregulation syndrome

Regardless of the mode of action, the abundance of effects and their distribution across the genome suggests that the mixing of the two parental genomes impacts multiple cellular processes. This conclusion is consistent with a recent study in rice showing a large number of loci involved in heterosis, whose identity is largely different in different crosses^26^. A common scheme that emerged from our genetic and phenotypic data is a dysregulation of key cellular functions in the hybrid. This includes (i) repression of respiration is largely alleviated, allowing cells to respire even when glucose is present (ii) size-dependent extension of the cell cycle is perturbed (iii) under conditions of ethanol stress, the hybrid maintains consistent growth, not showing the cell cycle slowdown observed in the parental strain.

The consistent heterotic phenotype of the hybrid is becoming more surprising when considering the large scale rewiring of the hybrid’s regulatory functions, which, naively, would be expected to perturb, rather than improve cellular functions. One possible explanation is that regulatory functions that function as safeguard mechanisms to limit biomass production or growth diverge more readily and are thereby more amendable to perturbations in the hybrid. This appears consistent with the phenotypes we observe: limiting respiration, when oxygen is available, may reduce the efficiency of carbon metabolism but can function to preserve resources and eventually to convert them to ethanol for later consumption and to defend against competing microorganisms. Prolonged G1 (or G2) increases division time, but allows regulation of the cell size, such as correcting for large fluctuations in size. Slowing down the cell cycle during ethanol stress may prevent damage, perhaps explaining why it was not reverted during evolution.

In this view, heterosis may be a reflection of the different tradeoffs that govern evolution: while growth rate needs to be maximized, this maximization is subject to some constraints, such as maintaining genome stability, a process that may indeed be limited in the hybrid, as we showed. Consistent with this hypothesis, several studies, including in plant hybrids, showed a tradeoff between growth and stress response^29^. Further, a role of evolutionary tradeoffs in constraining agriculture relevant traits such as biomass growth is well appreciated from studies of crop and animal breeding, selecting for plants for maximal agriculture productivity, for example, reduces their fitness under natural conditions.

Taken together, our study suggests that heterosis and incompatibility may be tightly linked: incompatibilities that alleviate safeguard mechanisms may contribute to hybrid growth vigor (Figure 7). Such an incompatibility-based explanation of heterosis may account for results that were not predicted from the dominance/complementation model, such as the maintenance of heterosis after deleterious mutations are purged, or progressive heterosis in polyploids^3^.

**Fig. 7.**
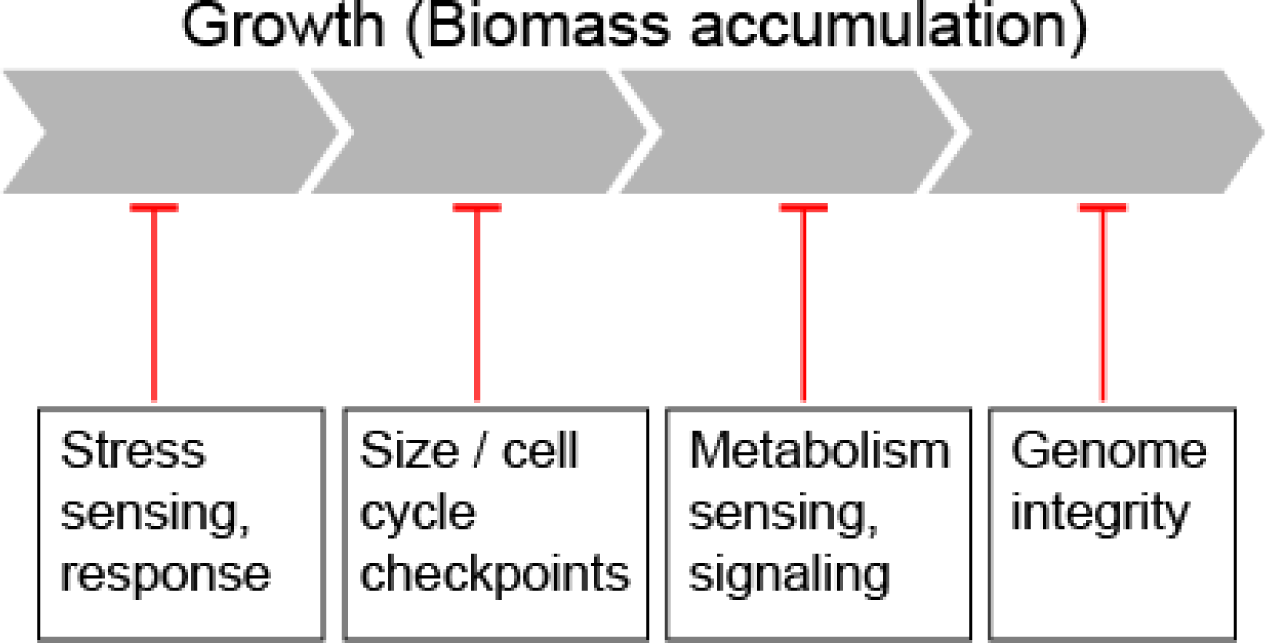
Model of heterosis: Hybrid incompatibilities perturbing failsafe mechanisms limit growth of the wild-type background.

## Materials and Methods

### Growth curves

The hybrid and the diploid parents were grown overnight in YPD and back-diluted to OD_600_ 0.05 in YPD and grown for 5 hours in 30° C, in order to allow the cultures to reach log phase. Cells were then diluted in the appropriate media, for growth in stress media-cells were washed twice in water previous to dilution. Cultures were grown at 30° C unless otherwise indicated. The cultures were kept in log phase by back diluting to OD_600_ 0.05 when a culture reached OD_600_ ~1.

### Cell cycle analysis using high throughput time-lapse microscopy

Olympus IX71 microscope was automated using motorized XY stage (Prior), fast laser autofocus attachment ^30^, excitation and emission filter wheels (Prior) and shutters (UniBliz). The EMCCD camera was AndoriXon with pixel size of 16 and 512x512 EMCCD chip cooled to -68C. eGFP and mCherry were detected using EXFO X-Cite 120 light source at 12.5% intensity using Chroma 89021 mCherry/GFP ET filter set. Exposure time for the detection of eGFP and mCherry was 100 msec. The cells were observed using 60x0.9 NA UPLFLN/APO objective. The microscope was controlled by custom written software running on Red Hat Linux. The fast auto focus and filter switching times allowed simultaneous imaging of 60 fields of view with time resolution of 3 minutes.

Preparation of cells for time lapse imaging was performed as previously described ^31^: Briefly, log stage cultures were seeded at OD_600_ 1 on a slab of 2% low melt agarose containing SC and imaged between the agar pad and the cover glass. Bright field images were taken 1 micron below the focal plane to facilitate image analysis. This time-lapse setup allows unperturbed exponential growth in a single plane throughout most of the experiment. Image analysis for the hybrid and *S. cerevisiae* was performed as previously described^32^, image analysis for *S. paradoxus* was performed by manually tracking the cell division.

### Respiration inhibition using Antimycin A

Three biological replicas of each strain were grown overnight in SD and back-diluted to OD_600_ 0.05 in 20 ml YPD. After allowing the cultures to reach log phase, they were washed in water and diluted to OD_600_ 0.01 in SD + 10μM Antimycin A media. The cultures were kept in log phase by back diluting to OD_600_ 0.01 when a culture reached OD_600_ ~1. OD_600_ measurements were taken every 90 minutes.

### Counting *RAD52* foci as indication of double strand breaks

Strains 4741 RAD52-YFP and CBS432 OS142 *S. paradoxus* Mata RAD52-YFP were constructed. Each of the strains was mated with Matα *S. paradoxus* and *S. cerevisiae*. In order to select diploid progenies following the mating, single colonies were collected and their DNA was stained in order to differ diploid colonies from haploids using flow cytometry. Once in log phase, cells were incubated in 70% ethanol, incubated in RNase A 1mg/ml for 40 minutes at 37°C, then incubated in Proteinase K 20 mg/ml for 1 hour incubation at 37°C, following 1 hour incubation in SYBR green (S9430, Sigma-Aldrich) (1:1000) in room temperature in the dark. Cells were washed twice with 50mM Tris-HCl pH8 between each incubation. The stained cells were sonicated in Diagenode bioruptor for 3 cycles of 10’’ ON and 20’’ OFF in low intensity. The fluorescence was measured using BD LSRII Flow Cytometer (BD Biosciences). After diploid colonies were picked, cells were grown in SD to log phase and were imaged in OD_600_ 0.150. Images were acquired using Olympus IX83 based Live-Imaging system equipped with CSU-W1 spinning disc: sCMOS digitale Scientific Grade Camera 4.2 MPixel VS LaserModule 1863C with LaserMerge System Laser module - laser 488nm with 100mW. The number of cells containing *RAD52* foci was counted.

### Mitochondrial morphology using electron microscopy

Cells were grown in SD medium. Once they reached an OD_600_ of ~0.5, 5ml of a fixative solution was added to 5ml media to a final concentration of 6% paraformaldehyde, 4% glutaraldehyde and Cacodylate buffer 0.2M (pH=7.4). The samples were gently rotated for 40 minutes at 30°C, then centrifuged and washed twice in Cacodylate buffer and incubated again for an hour in the fixative solution. After washing with Cacodylate buffer and centrifugation, samples were embedded in 10% gelatin in water, fixed over night with the fixative solution, washed in Cacodylate buffer and then incubated overnight in 2.3 M sucrose and rapidly frozen in liquid nitrogen. Frozen ultrathin (70-90 nm) sections were cut with a diamond knife (Diatome AG, Biel, Switzerland) at -120°C on an EM UC6 ultramicrotome (Leica Microsystems, Vienna, Austria). The sections were collected on 200-mesh formvar coated nickel grids. Contrasting and embedding were performed as previously described ^33^. The embedded sections were observed in a Tecnai T12 electron microscope (FEI, Eindhoven, The Netherlands) operating at 120 kV. Images were recorded using an ErlangshenES500W CCD camera (GATAN) or an Eagle 2k x2k CCD camera (FEI). The quantification of mitochondrial area and cristae was done using imageJ^34^.

### Cell Cycle imaging for time-lapse movie and stills

Cells were grown in SD to reach log phase. Samples with OD_600_ ~0.02 were seeded on a slab of SD + 2% slab low melt agarose. Images were taken in Olympus IX83 based Live-Imaging system equipped with CSU-W1 spinning disc: sCMOS digital Scientific Grade Camera 4.2 MPixel. The cells were kept at 30°C using in-stage incubator Chamlide TC. Images were taken every 5 minutes. Movie cropping and labeling was done using imageJ ^34^

### Measuring oxygen and glucose consumption

The fermentation experiments were performed using a DASBox mini Bacterial Fermentation system (DASGIP, Eppendorf), with the online monitoring and control of the temperature, dissolved oxygen (DO), OD, aeration and mixing. Yeast cells were grown in the fermenters in 200ml SC media supplemented with 2% glucose, using the following controlled parameters: 30°C, 300RPM, and 0.5 VVM of air. When oxygen became limiting (DO=<20%), a feedback cascade of mixing and aeration was engaged (300-800RPM, and 0.5-1.0VVM respectively). The runs were performed as follows: overnight starters were used and diluted into the fermenters to an OD_600_ of ~0.1 and grown as above. At the indicated time points, OD was measured also offline after a 5” sonication using the Sonics –VibraCel sonicator with a micro-tip (at 80% pmt) to break clamps of cells. Residual glucose levels were also measured offline using Accu-Chek Sensor strips (Roche).

### Reanalysis of gene expression data

Processed data was downloaded from Gene Expression Omnibus (GEO) with accession number GSE14708^13^. For each gene, a t-test was performed between the parental and hybrid gene expression, taking the three replicates together. Enrichment was tested using XL-mHG test on the ranked list of p-values (with parameters X=5,L=400).

### Proteomic analysis of diploids and hybrids GFP-tagged collections

*S. cerevisiae* 4742 Mata HO::Nat^r^ was systematically mated against the GFP collection (::HIS3; the library was a kind gift from J. Weissman, University of California, San Francisco, San Francisco, CA; ^35^). Mating was performed on rich media plates, and selection for diploid cells was performed on plates with clonNAT Nourseothricin (Werner) and lacking HIS. To manipulate the collection in high-density format (384), we used a RoToR bench top colony arrayer (Singer Instruments). Automatic high-throughput microscopy screens and analyses were performed as was described previously ^36^.

### Creating a hemizygote hybrid collection

The yeast MATa haploid deletion collection of non essential genes^37,38^ consisting of 5171 open reading frames replaced with G418 resistance was systematically mated with *S. paradoxus aNat* (Matα strain of *S. paradoxus* CBS432 OS142) to produce a collection of hemizygote hybrids. The strains were organized in 96 well plates (Nunc) and replicated onto YPD agar plates pre-plated with the *S. paradoxus* α strain. The plates were incubated over night at 30°C to allow mating to occur and then replicated to double selection plates (YPD + 0.1mg/ml Nat + 0.2mg/ml G418) to select for diploid progeny. The final collection consisted of 4484 hemizygote hybrid strains.

### Pooling

The hemizygote *S. cerevisiae* and hybrid libraries were grown on YPD agar plates with antibiotic selection of G418 (200 μg/ml, Calbiochem) for the hemizygote *S. cerevisiae* library and with G418 (200 μg/ml) and Nourseothricin (Nat, 200 μg/ml WERNER BioAgents) for the hybrid library. After two days of growth at 30°C, the libraries were replicated in triplicates. Following two days of growth at 30°C, colonies had a good and uniform size and cells were soaked off the plates by adding 10 ml of YPD liquid media + G418 (200 μg/ml) to each plate and gently scraping the cells off the plate to resuspend the cells. After the cells were resuspended, a fixed volume of 100 μl from each plate was removed with a pipette. These samples were all mixed together, diluted to have roughly an OD_600_ of 50 and kept in -80°C with a final concentration of 15% Glycerol. From the three copies for each library, the two best looking plates were taken and pooled in parallel; these represent the biological replicates we used for each library.

### Growth experiment of hemizygote libraries

The pools were thawed on ice. Then, for recovery, they were diluted to OD_600_ 0.1 in 30 ml of YPD and grew at 30°C for three hours, completing roughly 1.5-2 doublings. Subsequently, OD_600_ was measured, and the pools were diluted to OD_600_ 0.01 in a volume of 200ml. At this point, the first time sample was removed, depicting the reference time point measurement. Along the experiment, samples were taken every 3-4 generations, giving on average 9 time points over a total of 30 generations. Before the cells reached OD_600_ of 1, they were diluted to an OD_600_ of 0.01, in order to keep them in exponential phase throughout the experiment. For sampling, 6*10^7^ cells were removed twice (serving as technical replicates), spun down, supernatant removed and the pellet saved at -20°C. Given the size of the pools, every strain should be sampled ~10000 times, assuming equal representation, minimizing sampling biases.

### Genomic DNA purification and PCR amplification of barcodes

Genomic DNA was purified using Epicenter MasterPureTM Yeast DNA Purification Kit. Next, 100ng of genomic DNA was used for the PCR amplification, which was conducted in a total of 50 μl, using KAPA HiFi HotStart PCR Kit with the following conditions: 95°C/4 min; 23 cycles of 98°C/20 s, 65°C/15 s, 72°C/15 s; followed by 72°C/5 min. Barcodes were added to the general primers. For the upstream barcodes the following primers were used: (forward) 5′-NNNNNGATGTCCACGAGGTCTCT-3’ and (reverse) 5’-NNNNNGTCGACCTGCAGCGTACG-3’. For the downstream barcodes the following primers were used: (forward) 5’-NNNNNCGAGCTCGAATTCATCGAT-3’ and (reverse) 5’-NNNNNCGGTGTCGGTCTCGTAG-3’. PCR product was then purified with Qiagen MinElute 96 UF PCR Purification Kit. Following PCR purification, DNA was quantified with the Invitrogen Quant-iT dsDNA BR Assay Kit and then equal amounts of DNA were pooled.

### Sequencing analysis and growth rate quantification

The 5-base multiplexing tag allowed for post-sequencing assignment of each read to a particular measurement (time point and pool) using fastx_barcode_splitter allowing zero mismatches. Using cutadapt and fastx_trimmer all the multiplex barcodes and the common primer sequences were removed, to be left with the strain barcodes only. The strain barcodes were aligned to a reference table using bowtie allowing for one mismatch. All barcode counts (B1 for the barcode upstream of the selection marker and B2 for the barcode downstream of the selection marker) were separately normalized by total coverage (read counts were divided by total read count per time point and pool, multiplied by 10^6^ to get number of transcript per million (TPM)). For all conditions expect for YPD + 8% Ethanol, the PCR and sequencing was performed twice as mentioned above (serving as technical replicates). These technical repeats were averaged after total read normalization by taking the average in log scale. For each strain that had more than 30 reads at the first time point, growth rates were extracted by fitting a linear model (log2(TPM + 1) ~ generations) using the rlm function in R (MASS package) for B1 and B2 separately. The reported growth rate per strain is the mean of B1 and B2, if both B1 and B2 growth rates were defined, otherwise just the identified growth rate (B1 or B2). Under the growth condition of YPD + EtOH, only the first 5 time points were taken for calculating the growth rates, as the population size remained stable over this range. For each condition, growth rates were standard normalized (mean was subtracted and divided by standard deviation). A strain was considered to have reduced growth rate if the *Z* Score was below -1.5 in both biological replicates. Some strains could never be identified, presumably because of mutations in the barcode or the primer sequences. The number of strains that were identified in our screen in at least one condition was 4004 out of 4484 and 5362 out of 6330 for the hybrid and the *S. cerevisiae* pool respectively. The hybrid pool is significantly smaller to begin with because it excludes any essential strains.

### Growth assay in ethanol toxicity

Newly transformed 4741 MKS1::Hyg^r^, RTG2::Hyg^r^ and KanMX-pTDH3-RTG2 strains were mated with *S. paradoxus* Matα HO::Nat^r^ with the deletions MKS1::Hyg^r^ or RTG2::Hyg^r^ to create hybrids. After the mating they were grown on selection plates in order to select for diploid progenies. The strains were grown in YPD overnight in optimal conditions and back-diluted by adding 20μl of the stationary culture to 5ml YPD. After allowing the cultures to reach log phase, they were washed with YPD + 7.5% ethanol and diluted to OD_600_ 0.05 in YPD + 7.5% ethanol media. The cultures were back-diluted to OD_600_ 0.05 if during an OD_600_ measurement the culture OD_600_ was higher than 0.35.

## Acknowledgements

The EM studies were conducted at the Irving and Cherna Moskowitz Center for Nano and Bio-Nano Imaging (Weizmann Institute of Science). We thank members of our groups for discussions and Gilgi Friedlander for help with bioinformatics data processing. This work was supported by grants from the ERC, ISF, Minerva Center, ICORE and the Helmsley charitable trust.

## Author contributions

R.H.H., D.B., N.B. and A.L. designed the experiments and drafted the manuscript; R.H.H. performed and analyzed the pooled library screen experiments; D.B. and E.S. performed and analyzed the EM experiments. D.B. and G.Y. performed and analyzed the fermenter experiments. D.B, S.R. and I.S. performed and analyzed the cell cycle experiments. M.B. performed and analyzed the protein localization screen. M.S. discussed data interpretation; N.B. and A.L. supervised the study.

## References

1. Darwin, C. On the Origin of the Species. Darwin 5, (1859).

2. Darwin, C. The Effects of Cross- and Self-fertilization in the Vegetable Kingdom. John Murry (1876).

3. Birchler, J. A., Yao, H., Chudalayandi, S., Vaiman, D. & Veitia, R. A. Heterosis. Plant Cell 22, 2105–2112 (2010).

4. Landry, C. R., Hartl, D. L. & Ranz, J. M. Genome clashes in hybrids: insights from gene expression. Heredity (Edinb). 99, 483–93 (2007).

5. Mallet, J. Hybrid speciation. Nature 446, 279–283 (2007).

6. Schumer, M., Cui, R., Rosenthal, G. G. & Andolfatto, P. Reproductive Isolation of Hybrid Populations Driven by Genetic Incompatibilities. PLoS Genet. 11, (2015).

7. Mesgaran, M. B. et al. Hybridization can facilitate species invasions, even without enhancing local adaptation. Proc. Natl. Acad. Sci. U. S. A. 113, 10210–4 (2016).

8. Nolte, A. W. & Tautz, D. Understanding the onset of hybrid speciation. Trends Genet. 26, 54–58 (2010).

9. Duvick, D. N. Heterosis: feeding people and protecting natural resources. Genet. Exploit. heterosis Crop. 19–29 (1999).

10. Zu Ermgassen, E. K. H. J., Phalan, B., Green, R. E. & Balmford, A. Reducing the land use of EU pork production: where there’s swill, there’s a way. Food Policy 58, 35–48 (2016).

11. Steinmetz, L. M. et al. Dissecting the architecture of a quantitative trait locus in yeast. Nature 416, 326–330 (2002).

12. Lippman, Z. B. & Zamir, D. Heterosis: revisiting the magic. Trends Genet. 23, 60–66 (2007).

13. Tirosh, I., Reikhav, S., Levy, A. A. & Barkai, N. A yeast hybrid provides insight into the evolution of gene expression regulation. Science (80-.). 324, 659–662 (2009).

14. Bell, G. D. M., Kane, N. C., Rieseberg, L. H. & Adams, K. L. RNA-seq analysis of allele-specific expression, hybrid effects, and regulatory divergence in hybrids compared with their parents from natural populations. Genome Biol. Evol. 5, 1309–23 (2013).

15. Tirosh, I., Sigal, N. & Barkai, N. Divergence of nucleosome positioning between two closely related yeast species: genetic basis and functional consequences. Mol. Syst. Biol. 6, 365 (2010).

16. McManus, C. J., May, G. E., Spealman, P. & Shteyman, A. Ribosome profiling reveals post-transcriptional buffering of divergent gene expression in yeast. Genome Res. 24, 422–430 (2014).

17. Groszmann, M. et al. Changes in 24-nt siRNA levels in Arabidopsis hybrids suggest an epigenetic contribution to hybrid vigor. Proc. Natl. Acad. Sci. U. S. A. 108, 2617–22 (2011).

18. Shapira, R., Levy, T., Shaked, S., Fridman, E. & David, L. Extensive heterosis in growth of yeast hybrids is explained by a combination of genetic models. Heredity (Edinb). 113, 316–326 (2014).

19. Albertin, W. & Marullo, P. Polyploidy in fungi: evolution after whole-genome duplication. Proc. R. Soc. B Biol. Sci. 279, 2497–2509 (2012).

20. Kellis, M., Patterson, N., Endrizzi, M., Birren, B. & Lander, E. S. Sequencing and comparison of yeast species to identify genes and regulatory elements. Nature 423, 241–54 (2003).

21. Kron, S. J., Styles, C. a & Fink, G. R. Symmetric cell division in pseudohyphae of the yeast Saccharomyces cerevisiae. Mol. Biol. Cell 5, 1003–1022 (1994).

22. Lisby, M. & Rothstein, R. Choreography of recombination proteins during the DNA damage response. DNA Repair (Amst). 8, 1068–1076 (2009).

23. Smith, A. M. et al. Quantitative phenotyping via deep barcode sequencing. Genome Res. 19, 1836–1842 (2009).

24. Deutschbauer, A. M. et al. Mechanisms of haploinsufficiency revealed by genome-wide profiling in yeast. Genetics 169, 1915–1925 (2005).

25. Chae, S. et al. A Systems Approach for Decoding Mitochondrial Retrograde Signaling Pathways. Sci. Signal. 6, (2013).

26. Huang, X. et al. Genomic architecture of heterosis for yield traits in rice. Nature 537, 629–633 (2016).

27. Park, S. J. et al. Optimization of crop productivity in tomato using induced mutations in the florigen pathway. Nat. Genet. 46, 1337–42 (2014).

28. Jiang, K. et al. Tomato Yield Heterosis Is Triggered by a Dosage Sensitivity of the Florigen Pathway That Fine-Tunes Shoot Architecture. PLoS Genet. 9, e1004043 (2013).

29. Miller, M., Song, Q., Shi, X., Juenger, T. E. & Chen, Z. J. Natural variation in timing of stress-responsive gene expression predicts heterosis in intraspecific hybrids of Arabidopsis. Nat. Commun. 6, 7453 (2015).

30. Paran, Y. et al. High-throughput screening of cellular features using high-resolution light-microscopy; Application for profiling drug effects on cell adhesion. J. Struct. Biol. 158, 233–243 (2007).

31. Bean, J. M., Siggia, E. D. & Cross, F. R. Coherence and timing of cell cycle start examined at single-cell resolution. Mol. Cell 21, 3–14 (2006).

32. Soifer, I. & Barkai, N. Systematic identification of cell size regulators in budding yeast. Mol. Syst. Biol. 10, 761 (2014).

33. Tokuyasu, K. T. Application of cryoultramicroscopy to immunocytochemistry. J. Micros. 143, 139–149 (1986).

34. Abràmoff, M. D., Magalhães, P. J. & Ram, S. J. Image processing with imageJ. Biophotonics Int. 11, 36–41 (2004).

35. Huh, W. et al. Global analysis of protein localization in budding yeast. Nature 425, 686–691 (2003).

36. Breker, M., Gymrek, M. & Schuldiner, M. A novel single-cell screening platform reveals proteome plasticity during yeast stress responses. J. Cell Biol. 200, 839–850 (2013).

37. Winzeler, E. A. et al. Functional characterization of the S. cerevisiae genome by gene deletion and parallel analysis. Science 285, 901–6 (1999).

38. Giaever, G. et al. Functional profiling of the Saccharomyces cerevisiae genome. Nature 418, 387–391 (2002).

39. Jorgensen, P., Nishikawa, J. L., Breitkreutz, B. & Tyers, M. Systematic identification of pathways that couple cell growth and division in yeast. Science (80-.). 297, 395–400 (2002).

